# System x_c_^−^ imaging maps ferroptosis-linked redox remodeling in cancer

**DOI:** 10.64898/2026.06.03.729933

**Authors:** Oskar Vilhelmsson Timmermand, Abigail R. Barber, Madeleine E. George, Sofia N. dos Santos, Hannah E. Greenwood, Richard S. Edwards, Muhammet Tanc, Alejandro H. Uribe, Will E. Tyrrell, Jade Bowden, Rizwan Farooq, Oliver Maddocks, Neel Patel, Miguel M. Murillo, Jasper van der Aart, Timothy H Witney

## Abstract

Ferroptosis is a regulated non-apoptotic form of programmed cell death that is implicated in tumor suppression and the normal tissue damage response. While the link between redox stress and ferroptosis is well established, no non-invasive methods exist to assess ferroptosis *in vivo*. Here, we demonstrate that the redox-sensitive positron emission tomography radiotracer and system x_c_^−^substrate, ^18^F-(*S*)-4-(3-fluoropropyl)-L-glutamic acid ([^18^F]FSPG), serves as a non-invasive marker of tumor ferroptosis. Global changes in amino acids, glutathione, and system x_c_^−^ activity occurred before loss of membrane integrity in cells sensitive to ferroptosis, but not in resistant cells. Resistant cells sensitized to ferroptosis through nuclear factor erythroid 2-related factor 2 (NRF2) knockout had reduced glutathione and [^18^F]FSPG retention, which were rescued by ferroptosis inhibitors. *In vivo*, immune checkpoint blockade decreased ferroptosis-specific [^18^F]FSPG tumor retention prior to immune cell infiltration. Together, our data demonstrate that [^18^F]FSPG can identify early redox changes that precede ferroptosis and enabled real-time monitoring of immunotherapeutic efficacy.

## Introduction

Ferroptosis is an iron-mediated non-apoptotic form of programmed cell death (1) that is conserved across multiple kingdoms of life (2, 3). Preclinically, ferroptosis plays an essential role in tumor growth control and the response of cancer cells to numerous anti-cancer treatments, such as immunotherapy, chemotherapy, and radiotherapy (4–6). Moreover, ferroptosis is induced as part of normal tissue response to damage following ischemia (7), inflammation, and infection (reviewed by Chen et al (8)). Unfortunately, no appropriate non-invasive biomarker(s) of ferroptosis exist (9). This hinders the translation of current knowledge into the clinic, largely limiting research to observations made in isolated cells or tissue sections.

Key to this regulated form of cell death are phospholipid-poly unsaturated fatty acids (PL-PUFAs), which are incorporated into the phospholipid membranes of the cell by the enzyme long-chain-fatty-acid—CoA ligase 4 (ACSL4) (10, 11). These PL-PUFAs are the targets for reactive oxygen species (ROS), resulting in lipid peroxides that drive membrane disintegration and ferroptotic cell death. Iron-catalyzed conversion of peroxides to reactive peroxyl radicals propagates membrane destruction (12). Whilst the molecular basis of ferroptosis has not been fully elucidated, there are at least three major molecular pathways that regulate membrane peroxidation, including iron metabolism, the glutathione peroxidase 4 (GPX4)-system x_c_^−^ axis, and the ubiquinone (coenzyme Q10, coQ10) pathway (2, 13, 14).

The classical GPX4-system x_c_^−^ ferroptosis axis, first described in 2012 by Dixon *et al.* (1), identified GPX4 as the master regulator of ferroptosis, acting to neutralize deleterious lipid peroxides by converting them into lipid alcohols (15). GPX4 activity requires glutathione (GSH), an important cellular antioxidant. GSH is a tripeptide, synthesized from glutamate, cysteine and glycine; cysteine availability being limiting for its synthesis (16). In tumors, cysteine is predominantly supplied through the import of its oxidized form, cystine, by the cystine/glutamate antiporter, system x_c_^−^. System x_c_^−^ is a heterodimer, consisting of the light chain transporter, xCT, and a heavy chain, 4F2hc, which is required for membrane localization. Inhibition of system x_c_^−^, for example, by the small molecule imidazole ketone erastin, results in the depletion of intracellular pools of GSH and induction of ferroptosis (17). Conversely, the metabolic reprogramming of cancer frequently results in upregulation of both GSH and system x_c_^−^ (18), which may, in turn, confer resistance to ferroptosis (19).

We have previously used ^18^F-(*S*)-4-(3-fluoropropyl)-L-glutamic acid ([^18^F]FSPG), a fluorine-18 labelled glutamate analog, to non-invasively determine system x_c_^−^ activity, assess treatment response, and predict resistance to chemotherapy using positron emission tomography (PET) (18, 20–22). Given the direct link between GSH, redox dysregulation and the ferroptotic response, we hypothesized that [^18^F]FSPG PET imaging would provide an early marker of ferroptosis *in vivo*. Using a range of cellular, pharmacological, and genetic models across a range of therapies, we show that system x_c_^−^ activity and associated [^18^F]FSPG retention is a sensitive marker of ferroptosis-associated redox remodeling that can be used to predict the efficacy of checkpoint inhibitors.

## Results

### Ferroptosis accompanies global changes in GSH metabolism and [^18^F]FSPG retention

To establish the link between [^18^F]FSPG tumor retention and the GPX4-system x_c_^−^ axis, we treated several cancer cell lines grown in culture with varying concentrations of RSL3, a small molecule drug that inhibits GPX4 and induces ferroptosis (14, 23). These cells had a range of sensitivities to GPX4 inhibition, with cell death rescued with co-treatment with the ferroptotic inhibitor and membrane ROS scavenger, liproxstatin-1 (**Supplementary Fig 1a**). From our panel of cells, we selected the human sarcoma cell line HT1080, a cell line sensitive to RSL3, and a non-small cell lung cancer cell line (A549) that was resistant to RSL3 treatment for further evaluation.

xCT is an integral part of system x_c_^−^ and comprises 12 transmembrane helices required for bidirectional cystine/glutamate transport (24). In both HT1080 and A549 cells, xCT protein expression was increased in a dose-dependent manner following RSL3 treatment. The magnitude of these changes was far greater in HT1080 cells (**Fig. 1a**). Besides xCT, the iron-storage protein ferritin (FTH1) was also upregulated in both cell lines at higher concentrations of RSL3. Other proteins known to mediate ferroptosis, including nuclear factor erythroid 2-related factor 2 (NRF2), acyl-CoA synthetase long chain family member 4 (ACSL4), GPX4, and nuclear receptor coactivator 4 (NCOA4) were either unchanged or decreased. ACSL4, GPX4 and NCOA4 were reduced in HT1080 cells with RSL3, whereas these were not altered in A549 cells. Lipid peroxidation and cell necrosis increased in HT1080 cells over a range of RSL3 concentrations, which did not occur in the more resistant A549 cells (**Fig. 1b, c**).

**Figure 1.**
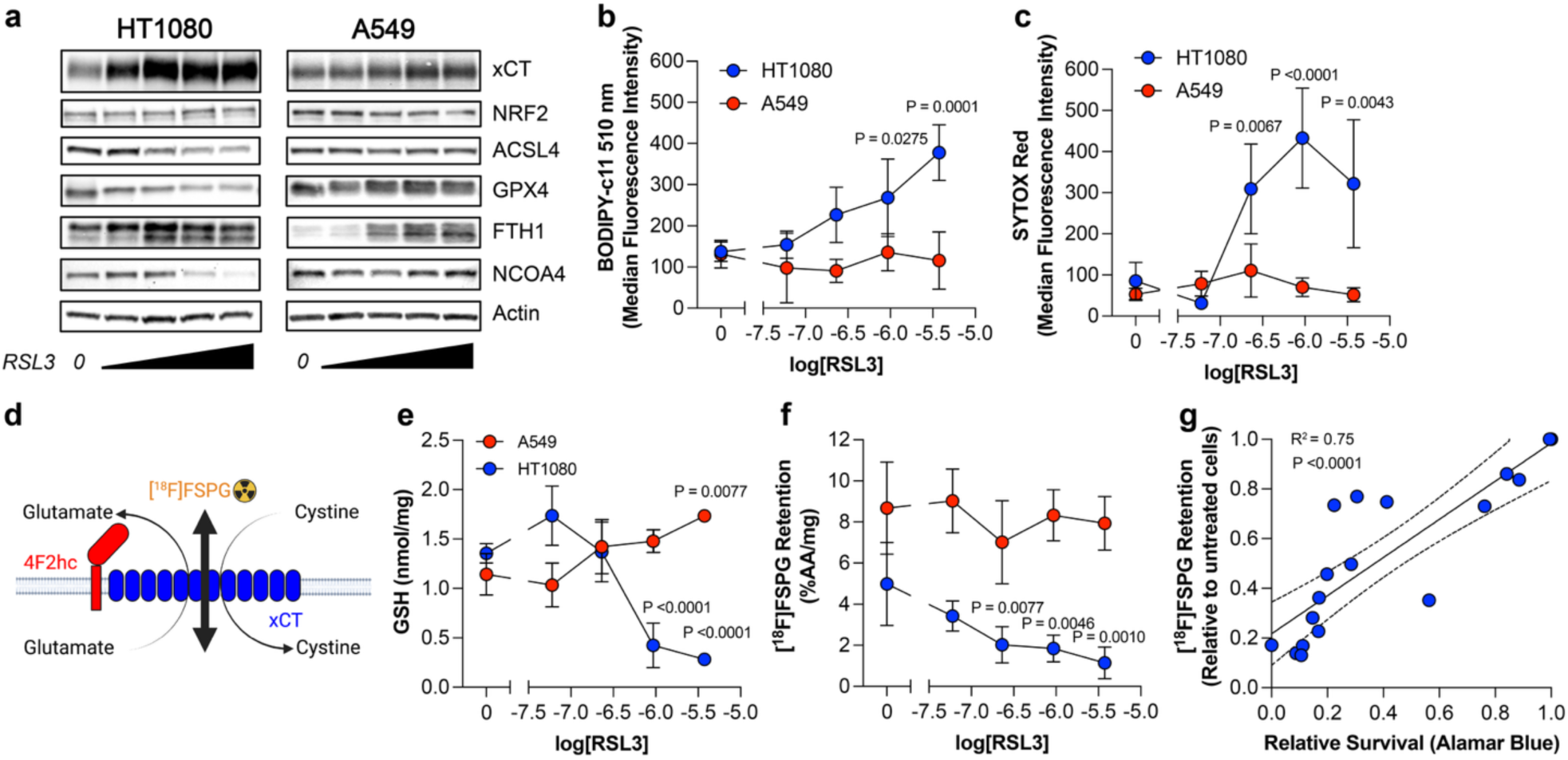
Ferroptosis alters GSH metabolism and reduces [^18^F]FSPG retention in sensitive tumor cells. **a,** Western blot of xCT, NRF2, ACSL4, GPX4, FTH1, NCOA4 and actin protein expression in HT1080 and A549 cell lines treated with increasing concentrations of RSL3 (0, 0.06, 0.23, 0.93, 3.75 µM). **b,** Lipid peroxidation, measured with BODIPY C11. **c,** Cell necrosis, measured with Sytox Red. **d,** Scheme depicting system x_c_^−^ function and [^18^F]FSPG retention. **e,** Total GSH following RSL3 treatment. **f,** RSL3 dose-dependent changes in [^18^F]FSPG retention in HT1080 and A549 cells. %AA, percent added radioactivity per mg protein. **g**. Linear correlation between [^18^F]FSPG retention and relative survival measured with Alamar blue in HT1080 cells. Dotted lines represent the 95% confidence interval.

Given the profound changes in xCT expression that occurred during ferroptosis, we next asked whether [^18^F]FSPG retention might provide a non-invasive marker of this important pathway. [^18^F]FSPG is bidirectionally transported across the plasma membrane by system x_c_^−^ (**Fig. 1d**), where its retention is inversely proportional to the rate of *de novo* GSH biosynthesis (20). GSH was depleted in HT1080 cells treated with RSL3, while being maintained in A549 cells (**Fig. 1e)**. Importantly, retention of [^18^F]FSPG decreased in a dose-dependent manner following RSL3 treatment in HT1080 cells, whereas it was preserved in A549 cells (**Fig. 1f**). These changes in [^18^F]FSPG retention correlated to survival in HT1080 cells (**Fig. 1g**) but not with total GSH, which was only depleted at high concentrations of RSL3 (**Supplementary Fig. 1b**) and remained unchanged in A549 cells (**Supplementary Fig. 1c**). This pattern was not exclusive to human cells. Similarly to HT1080 cells, GPX4 inhibition resulted in a dose-dependent decrease in [^18^F]FSPG in a mouse melanoma cell line (B16-F0), which was sensitive to ferroptosis (**Supplementary Fig. 1d**).

To get a better understanding of the mechanisms driving the changes in both xCT and GSH, we quantified intracellular steady-state amino acid concentrations using metabolomics. Whilst RSL3 treatment reduced global amino acid concentrations in HT1080 cells compared to untreated controls, these were increased in A549 cells; changes that were almost completely restored with the ferroptosis inhibitors ferrostatin-1 and liproxstatin-1 in both cell lines (**Fig. 2a**). A notable exception to this pattern was the intracellular concentrations of cystine, which was raised in HT1080 cells treated with RSL3 – highlighting the importance of system x_c_^−^ in cells undergoing ferroptosis following GPX4 inhibition. Whilst ferrostatin-1 and liproxstatin-1 treatments restored metabolomic measurements to baseline following GPX4 inhibition, xCT remained upregulated (**Fig. 2b**). [^18^F]FSPG retention was rescued in HT1080 cells treated with RSL3 in combination with ferroptosis inhibitors liproxstatin-1 and ferrostatin-1, whereas no changes were seen in the resistant A549 cells, directly linking radiotracer retention to ferroptotic cell death (**Fig. 2c**). Normalization of [^18^F]FSPG cell retention following dual RSL3 and ferroptosis inhibitor treatment in HT1080 cells was accompanied by restoration of lipid peroxidation (**Fig. 2d**), GSH (**Fig. 2e**), and necrosis to baseline levels (**Supplementary Fig. 2a**). No changes were observed in A549 cells under any of the treatment conditions. Changes in [^18^F]FSPG were also rescued by liproxsatin-1 but not ferrostatin-1 in B16-F0 cells (**Supplementary Fig. 2b**). Finally, to ascertain whether decreased [^18^F]FSPG retention was ferroptosis-specific, we incubated HT1080 cells with RSL3 in combination with inhibitors of ferroptosis, apoptosis, autophagy, and necroptosis. RSL3 decreased [^18^F]FSPG retention and was only rescued in cells treated with the ferroptosis inhibitor (**Fig. 2f**). Taken together, ferroptosis elicits large changes in [^18^F]FSPG cellular retention, which is accompanied by elevated xCT expression, GSH consumption, and the uptake of precursors required for *de novo* GSH biosynthesis.

**Figure 2.**
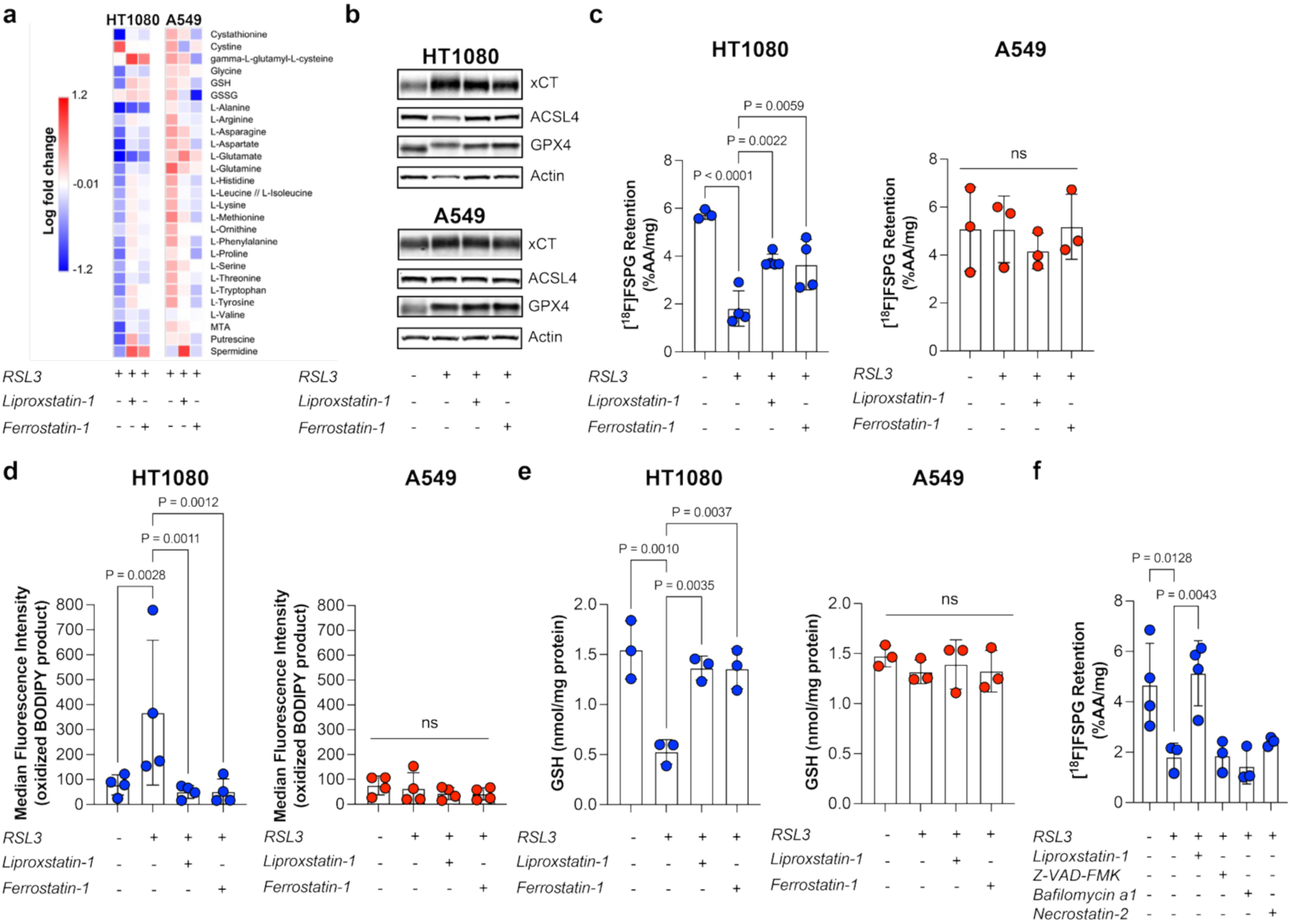
Ferroptosis inhibition restores [^18^F]FSPG retention in tumor cells. **a,** Amino acid metabolomics of HT1080 and A549 cells following RSL3 treatment (0.93 µM) with/without co-treatment with liproxstatin-1 (100 nM) or ferrostatin-1 (12 µM) compared to untreated cells (DMSO). **b,** Western blot of xCT, ACSL4, GPX4, and actin, protein expression in HT1080 and A549 cell lines treated with RSL3 (0.93 µM) alone and with the addition of liproxstatin-1 or ferrostatin-1. **c,** [^18^F]FSPG retention in HT1080 and A549 cells following RSL3 (0.93 µM) alone and with the addition of liproxstatin-1 or ferrostatin-1. **d,** Lipid peroxidation, measured with BODIPY C11 in HT1080 and A549 cells following RSL3 (0.93 µM) alone and with the addition of liproxstatin-1 or ferrostatin-1. **e,** Total GSH in HT1080 and A549 cells following treatment with RSL3 (0.93 µM) alone and with the addition of liproxstatin-1 or ferrostatin-1. **f,** [^18^F]FSPG retention in HT1080 cells following RSL3 treatment (0.93 µM) and treatment with cell death inhibitors of ferroptosis (liproxstatin-1, 100 nM), apoptosis (Z-VAD-FMK, 20 µM), autophagy (bafilomycin a1, 1µM), and necroptosis (necrostatin-2, 2 µM). ns, not significant.

### Redox and [^18^F]FSPG changes precede membrane rupture following induction of ferroptosis

Given the ability of [^18^F]FSPG to measure ferroptotic cell death in cells, we next asked whether a reduction in radiotracer retention reflected early biochemical changes that preceded ferroptosis, or whether they coincided with loss of membrane integrity. Ferroptosis was induced in HT1080 and the murine ovarian cancer ID8 cells, which both had similar sensitivities to RSL3 (**Supplementary Fig. 1a**). Just one hour after RSL3 treatment, [^18^F]FSPG retention was halved in these cells (**Fig. 3a, b**), a time at which xCT expression was also increased (**Fig. 3a, b** *inset*), and, or be it variably, lipid peroxidation induced (**Supplementary Fig. 3a**). Importantly, membrane integrity was still intact at this time (**Supplementary Fig. 3b**). In both cell lines, [^18^F]FSPG retention decreased during the first 3 h and then remained suppressed (**Fig. 3a, b**). The nadir at 3 h coincided with a peak in the necrotic phenotype and lipid peroxidation. The expression of xCT increased over the first six hours of treatment. In contrast, no such change was seen in GPX4 or ACSL4 (**Supplementary Fig. 3c**). Similarly to [^18^F]FSPG retention, a time-dependent decrease in total GSH concentration was measured, which was restored by 6 h in HT1080 cells, but remained low in ID8 cells (**Fig. 3c, d**, respectively).

**Figure 3.**
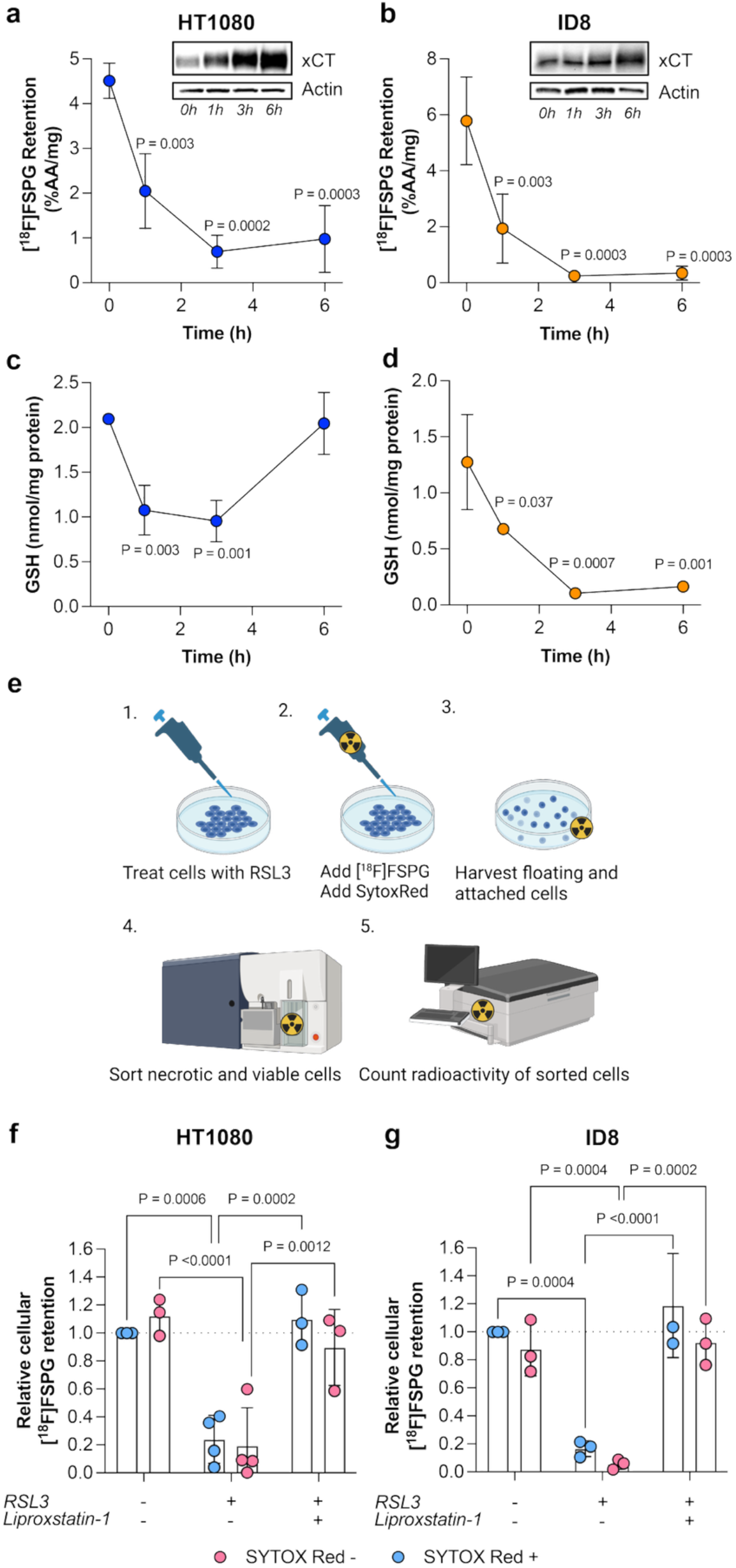
Early temporal changes in [^18^F]FSPG are accompanied by alterations in GSH metabolism. **a-b,** [^18^F]FSPG retention and protein expression of xCT (inset), during the first 6 h of RSL3 treatment (3.75 µM) in HT1080 cells (**a**) and in ID8 cells (**b**) following RSL3 treatment. **c-d,** Total GSH in HT1080 (**c**) and ID8 cells (**d**) during the first six hours of RSL3 treatment (3.75 µM)**. e,** Scheme describing the sorting of RSL3 cells co-incubated with Sytox Red and [^18^F]FSPG. **f-g,** [^18^F]FSPG retention in Sytox Red-positive and-negative cells relative to untreated viable controls in HT1080 (**f**) and ID8 (**g**) cells. Cells were treated with RSL3 (3.75 µM) for 3 h with and without liproxstatin-1 (100 nM) or remained untreated.

To completely exclude membrane disintegration as the driver of decreased [^18^F]FSPG retention, HT1080 and ID8 cells treated with RSL3 for 3 h were co-incubated with [^18^F]FSPG and the membrane impermeable dye, Sytox Red. Sytox Red binds DNA only once membrane integrity is compromised, providing a readout of cellular necrosis. Sytox Red positive and negative cells were sorted by fluorescence-activated cell sorting to separate necrotic from viable cells, with [^18^F]FSPG retention subsequently assessed in both populations by gamma counting (**Fig. 3e**). In untreated cells, [^18^F]FSPG retention remained high in both viable and necrotic cells, whereas [^18^F]FSPG was decreased in all populations following RSL3 treatment (**Fig. 3f, g**). This decrease in [^18^F]FSPG was restored in viable and necrotic cells upon liproxstatin-1 co-treatment. Our data suggest that [^18^F]FSPG traces early alterations in GSH metabolism that accompany ferroptosis. These changes in [^18^F]FSPG precede, and are not a consequence of, membrane permeabilization.

### NRF2 knockout and inhibition of FSP1 sensitize resistant cells to ferroptosis, which is detectable by [^18^F]FSPG

As we showed above, GPX4 inhibition fails to induce ferroptosis in A549 cells. A549 cells have aberrant activation of nuclear factor erythroid 2-related factor 2 (NRF2) (18), which controls the transcription of >200 antioxidant and cytoprotective genes through binding to antioxidant response elements (25). Under normal conditions, NRF2’s negative regulator, kelch-like ECH-associated protein 1 (KEAP1), tightly controls NRF2 via proteasomal degradation (26). A549 cells, however, have a mutation in KEAP1 (G333C), which prevents NRF2 degradation, enabling its translocation to the nucleus and activation of the cell’s antioxidant response. NRF2 has been suggested as the central controller of a network of different ferroptotic pathways – among them the genes for xCT (*SLC7A11*), GPX4 (*MCSP*), and ferroptosis suppressor protein 1 (FSP1, *AMID*) (27).

We have previously shown that [^18^F]FSPG is a sensitive marker of NRF2 pathway activation (18). To study the role of NRF2 in ferroptosis resistance, we treated A549 cells lacking NRF2 (A549 KO) (28) with RSL3. Unlike in parental A549 cells (**Fig. 1a**), there was a marked upregulation of xCT in A549 KO cells following GPX4 inhibition, which was restored with ferroptosis inhibitors liproxstatin-1 and ferrostatin-1. Additionally, three ferroptosis-associated proteins: ACSL4, FTH1, and NCOA4, were downregulated in comparison to untreated and rescued cells, whereas GPX4 levels were unchanged (**Fig. 4a**). The changes in xCT and GPX4 expression in response to RSL3 in A549 KO cells were similar to the response observed in the RSL3-sensitive HT1080 cells. Under the same treatment conditions, lipid peroxidation was significantly increased in A549 KO cells (**Fig. 4b**). In contrast, these ferroptosis markers were unchanged in parental cells (**Fig. 2d**). In A549 KO cells, RSL3 treatment resulted in loss of membrane integrity (**Fig. 4c**) and a corresponding decrease in GSH (**Fig. 4d**). These changes were detectable with [^18^F]FSPG, whose retention was halved following RSL3 treatment of A549 KO cells (**Fig. 4e**). All markers of ferroptosis, GSH metabolism, and [^18^F]FSPG were successfully rescued with liproxstatin-1 or ferrostatin-1 co-treatment. Restoration of functional NRF2 in A549 KO cells through ectopic expression rescued the wild-type phenotype, preventing RSL3-induced lipid peroxidation, necrosis, and changes in [^18^F]FSPG retention (**Supplementary Fig. 4a-c**).

**Figure 4.**
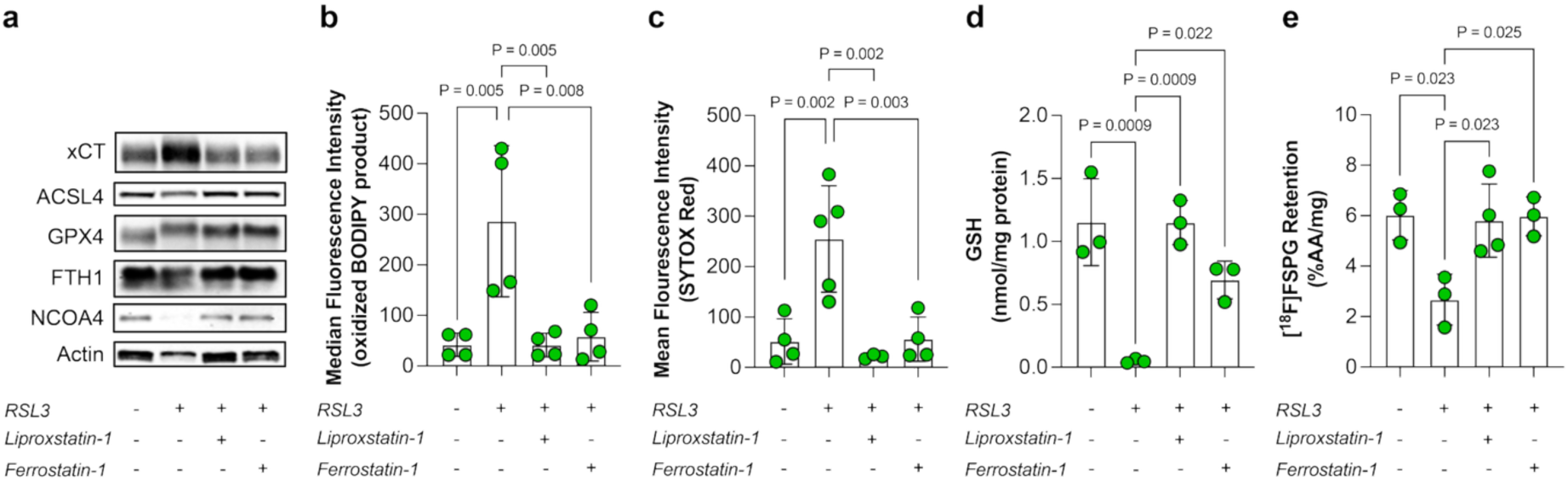
Loss of NRF2 sensitizes resistant cells to ferroptosis, which can be measured with [^18^F]FSPG. **a,** Protein expression of xCT, ACSL4, GPX4, FTH1 and NCOA4 in A549 NRF2 knockout cells (A549 KO). **b,** Lipid peroxidation in A549 KOs. **c,** Necrosis, measured using SYTOX Red, in A549 KOs treated with RSL3 (0.93 µM) and ferroptosis inhibitors (24 h). **d,** Total GSH in A549 KOs treated with RSL3 (0.93 µM) alone or in combination with liproxstatin-1 or ferrostatin-1 (100 nM and 12 µM respectively). **e,** [^18^F]FSPG retention in A549 KOs, treated with RSL3 (0.93 µM) and ferroptosis inhibitors (24 h).

Reduced coenzyme Q10 (ubiquinol) is another key ferroptosis surveillance pathway that complements the system x_c_^−^-GPX4 axis. Ubiquinol scavenges lipid peroxyl to generate ubiquinone. This reaction is catalyzed by FSP1, also known as anti-apoptosis-inducing factor mitochondria-associated 2 (AIFM2), which protects against ferroptosis (13, 29). As NRF2 transcriptionally regulates FSP1 (30), linking this axis to GSH metabolism, we next sought to understand whether inhibition of FSP1 (iFSP1) potentiated RSL3 treatment and whether [^18^F]FSPG provided a surrogate readout of this response. Combined RSL3 and iFSP1 upregulated xCT expression in both A549 and HT1080 cells, whilst dramatically reducing FSP1 expression (**Fig. 5a**). Dual treatment also reduced GSH (**Fig. 5b**) and [^18^F]FSPG retention in both cell lines (**Fig. 5c**), which was not seen in A549 cells for single agent RSL3 and was not fully rescued by 100 nM liproxstatin-1 (**Fig. 5c**). Lipid peroxidation was significantly increased in HT1080 cells treated with iFSP1 and RSL3 (**Supplementary Fig. 5**), but not in HT1080 or A549 cells treated with RSL3 alone. In sum, loss of NRF2 or inhibition of FSP1 sensitized resistant cells to ferroptosis, the magnitude of which was measurable with [^18^F]FSPG.

**Figure 5.**
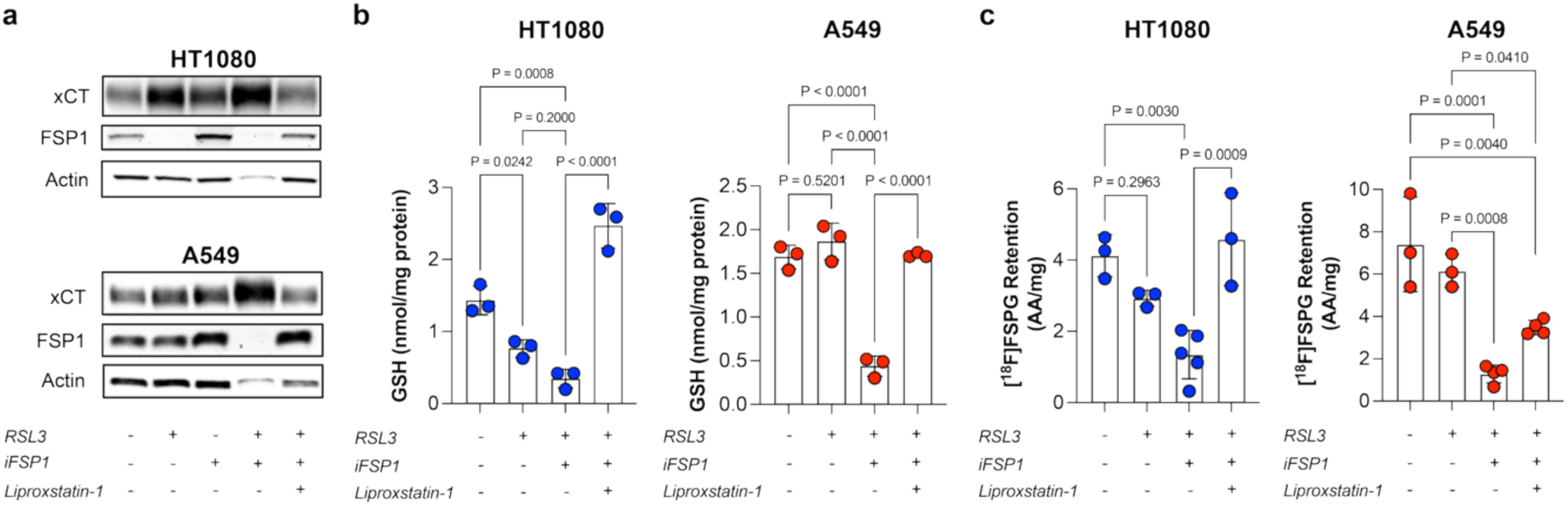
Ferroptosis following FSP1 inhibition can be measured with [^18^F]FSPG. **a,** xCT and FSP-1 protein expression in HT1080 and A549 cells challenged with RSL3 (0.06 and 0.93 µM respectively), iFSP1 (3 µM), and liproxstatin-1 (100 nM). **b,** Total GSH in HT1080 and A549 cells treated with RSL3 (0.06 and 0.93 µM respectively), iFSP1 (3 µM), and liproxstatin-1 (100 nM). **c,** [^18^F]FSPG retention in HT1080 cells and A549 cells challenged with RSL3 (0.06 and 0.93 µM respectively), iFSP1 (3 µM), and liproxstatin-1 (100 nM).

### [^18^F]FSPG is an early indicator of checkpoint blockade efficacy

Combination treatment with the checkpoint inhibitors anti-CTLA4 and anti-PDL1 is known to work in part by T-cell-mediated ferroptosis (4). Given that [^18^F]FSPG is robustly decreased following the induction of ferroptosis *in vitro*, we reasoned that [^18^F]FSPG imaging may provide an early readout of response to immunotherapy *in vivo*. To test this hypothesis, we treated the murine melanoma cell line B16-F0, a model known to respond well to checkpoint blockade, with anti-CTLA4 and anti-PDL1. [^18^F]FSPG PET imaging was characterized by high retention in untreated B16-F0 tumor xenografts, with non-tumor-associated radioactivity rapidly cleared by the kidneys into the bladder (**Fig. 6a**). [^18^F]FSPG retention in tumors was reduced by ∼40% at both three and seven days following immune checkpoint inhibition (ICI; 3.2 ± 0.25% ID/g and 3.0 ± 1.0% ID/g, respectively) compared to untreated (5.2 ± 1.3 %ID/g) and isotype control-treated mice (5.2 ± 1.0 %ID/g; **Fig. 6a, b**). Importantly, [^18^F]FSPG tumor retention was restored in combined liproxstatin-1 and ICI-treated mice (5.6 ± 1.3% ID/g), linking alterations in redox homeostasis and ferroptosis *in vivo*, which was detectable non-invasively with [^18^F]FSPG PET (**Fig. 6b**).

**Figure 6.**
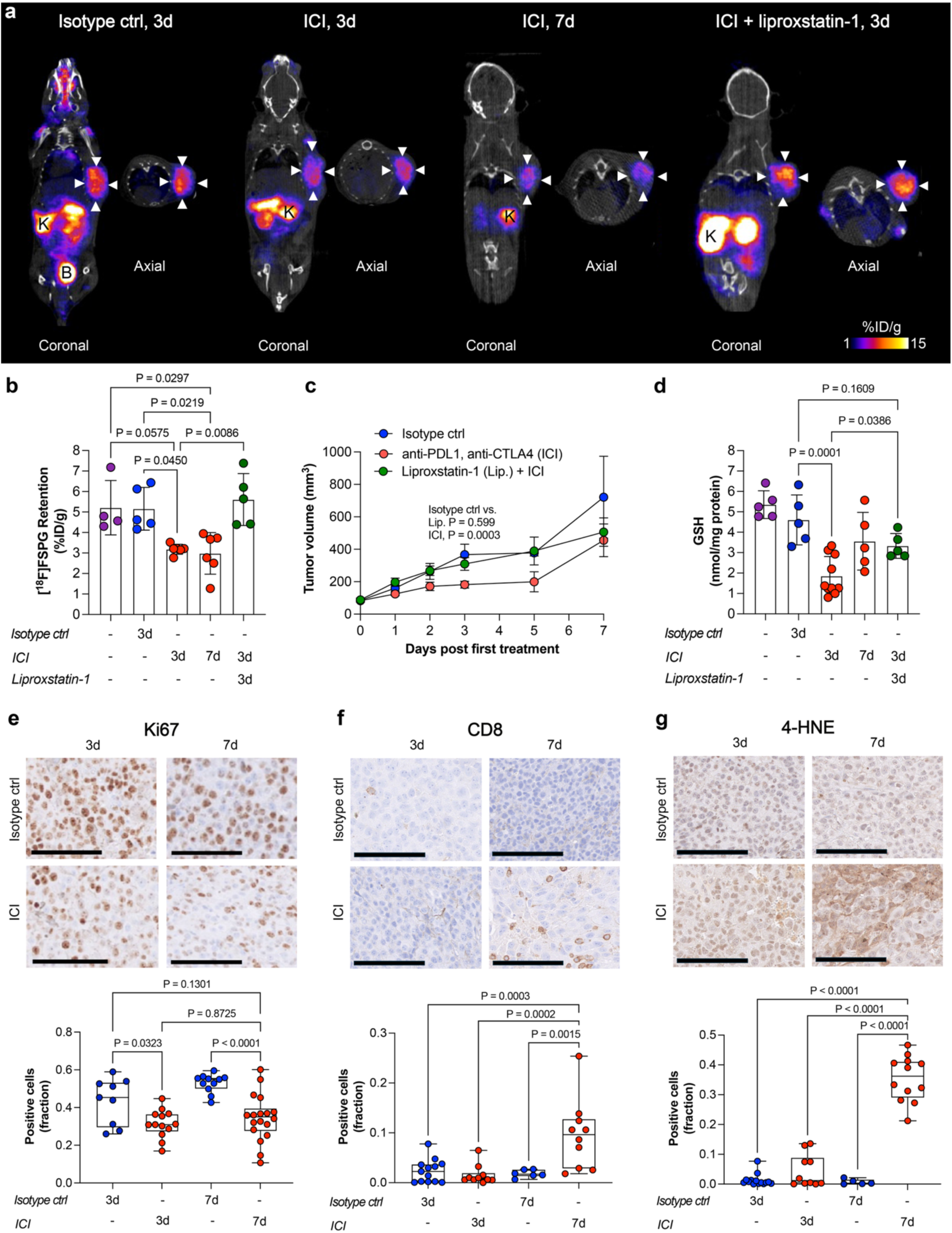
Imaging immune-mediated ferroptosis with [^18^F]FSPG PET. **a,** Representative coronal and axial [^18^F]FSPG PET/CT images in mice bearing B16-F0 xenografts before and after treatment with immune checkpoint inhibitors (ICI) or an isotype control (ctrl) antibody. ICI was additionally combined with liproxstatin-1. White arrows outline the tumor. K, kidney; B, bladder. **b,** Quantified [^18^F]FSPG retention in B16-F0 xenografts before and following ICI treatment, with and without liproxstatin-1. **c,** B16-F0 xenograft tumor volumes from mice treated with ICI as above, anti-PD-L1 (100 µg) and anti-CTLA4 (100 µg), dual isotype control (2 × 100 µg), or anti-PD-L1 and anti-CTLA4 together with daily liproxstatin-1. ICI, immune checkpoint inhibitor treatment. **d,** Total ex vivo GSH in B16-F0 xenografts following checkpoint blockade, isotype control or checkpoint blockade combined with daily liproxstatin-1 treatment. **e-g,** Representative immunohistochemical images and corresponding quantification of Ki67 (**c**), CD8 (**d**) and 4-HNE (**e**) staining in tissue sections from B16-F0 xenografts treated with ICI or isotype control antibody. The tissue was taken 3 or 7 days after the start of treatment. Scale bar, 100 µm.

As expected, *in vivo* growth of B16-F0 xenografts in immunocompetent mice was slowed by anti-CTLA4 and anti-PDL1 – an effect that was blunted with liproxstatin-1 (**Fig. 6c**). Three days after initiation of ICI treatment, a time in which tumor [^18^F]FSPG was significantly reduced, GSH was decreased by ∼65% compared to both pre-treatment controls and mice treated with an isotype control antibody (**Fig. 6d**). Partial but significant rescue of GSH was observed with liproxstatin-1 co-treatment, suggesting that early immune-mediated changes were connected to the ferroptotic phenotype. Ki67 – a proliferation marker – was lower in ICI-treated xenografts at both three and seven days (**Fig 6e**), indicating therapeutic efficacy and coinciding with changes in GSH and [^18^F]FSPG tumor retention. Interestingly, redox remodeling and decreased tumor cell proliferation occurred prior to significant immune cell infiltration, which occurred seven days after the start of treatment (**Fig. 6f, Supplementary Fig. 6a,b**) and corresponded with a significant increase in lipid peroxidation (**Fig. 6e**). Cleaved caspase-3, a marker of apoptosis, was not observed in any treatment condition (**Supplementary Fig. 6c**). Positive staining controls are displayed in **Supplementary Fig. 6d**.

Taken together, ICI treatment induces profound changes in immune cell composition in the tumor microenvironment, culminating in increased immune cell infiltration and lipid peroxidation by seven days. These changes are, however, preceded by alterations in redox status, tumor cell proliferation, and [^18^F]FSPG tumor retention, which are observable just three days after initiation of treatment.

## Discussion

The discovery of ferroptosis and other forms of programmed necrosis has fundamentally changed our view of programmed cell death. Loss of membrane integrity and intracellular content is inflammatory and, therefore, was viewed as an undesirable cell fate – one that the cell had no control over. This view changed through the pioneering work that identified several mediators of non-apoptotic cell death, such as GPX4 and FSP1 (1, 13). Mechanistic studies of these pathways, however, have been largely confined to investigations in isolated cells; until now, non-invasive assessment of ferroptosis in living subjects has not been possible. Using genetic, pharmacologic, and immunotherapeutic methods, we show here that the PET radiotracer [^18^F]FSPG provides an early and sensitive readout of ferroptosis-associated redox remodeling *in vivo*. The ability to assess ferroptosis in animal models, and in the future, humans, has widespread implications for monitoring therapy response, following the etiology of disease, and understanding the role of ferroptosis in normal physiology.

Whilst methods to evaluate ferroptosis *in vitro* are well established, definitive assessment of ferroptosis *in vivo* is challenging due to the low stability of many ferroptosis-inducing compounds and inhibitors, imperfect animal models, and deficiency of ferroptosis-specific biomarkers(31). Despite this, initiatives to visualize ferroptosis *in vivo* have been developed, including dyes that detect both apoptosis and ferroptosis (32) and magnetic resonance imaging agents for acute cardiac and kidney injuries (33). These methods, however, lack specificity and rely on techniques for detection that have poor tissue penetration and low sensitivity, respectively. A classical method to induce ferroptosis is to inhibit the activity of system x_c_^−^. We, and others, have shown that this inhibition, using erastin (and its derivatives) or sulfasalazine, can be non-invasively imaged by system x_c_^−^ radiotracers (22, 34–36). However, system x_c_^−^ inhibitors block cell uptake of these specific radiotracers independent of ferroptotic induction, making it an unreliable readout. With the emergence of multiple novel ferroptosis-inducing agents with potential *in vivo* applicability (37), techniques that can be used in preclinical and clinical settings to study and spatiotemporally track ferroptosis are urgently needed.

Redox plays an integral role in ferroptosis. Through the system x_c_^−^-GPX4 axis, GSH biosynthesis and recycling govern whether cells survive lipid peroxidation and subsequent loss of membrane integrity during ferroptosis(36). In our models, GSH was rapidly depleted in cells following the initiation of the ferroptotic cascade, halving just one hour after GPX4 inhibition (**Fig. 3**). These redox changes coincided with increased xCT expression, consequently elevating cellular concentrations of cysteine for *de novo* GSH biosynthesis (**Fig. 2**). Redox changes were absent in cells resistant to ferroptosis when treated with RSL3. Through FSP1 inhibition and knockout of NRF2, a direct regulator of GSH biosynthesis, resistant cells were sensitized to ferroptosis and GSH reduced – directly linking antioxidant production to cell fate. [^18^F]FSPG tracked these redox-related changes: cell retention of the radiotracer was extensively reduced at an early stage in cells undergoing ferroptosis, the reduction in [^18^F]FSPG was rescued only by ferroptotic inhibitors and not inhibitors of other cell death pathways, and no changes in [^18^F]FSPG were measured in resistant cells. We have previously shown that cystine directly exchanges with [^18^F]FSPG, resulting in reduced [^18^F]FSPG cell retention when rates of *de novo* GSH synthesis are increased (20), providing a possible mechanism behind ferroptosis-associated decrease in [^18^F]FSPG.

Through the imaging of ferroptosis we have uncovered new insights into the spatiotemporal dynamics of tumor response to immunotherapy. Ferroptosis is associated with immunotherapy efficacy (4, 38), but the importance of this mechanism and the timing of ferroptotic cell death following treatment remains unknown. Here, changes in redox homeostasis were an early marker of immune checkpoint blockade in B16-F0 murine tumors grown in immunocompetent mice, which preceded immune cell influx into the tumor and consequent lipid peroxidation (**Fig. 6**). This is important, as checkpoint inhibition has variable efficacy in solid tumors even when biopsy samples have high expression of the respective target (39). An early marker of treatment response will enable transfer of the patient to alternative second-line therapies with the opportunity to improve outcomes and reduce costs to the healthcare system. Current gold-standard imaging methods rely on imaging immune cell infiltration (40), which in tumor-bearing mice occurred many days after the initial redox-related perturbations, which were detected by [^18^F]FSPG PET.

One of the few limitations of using [^18^F]FSPG PET for ferroptosis imaging is the need for a baseline measurement, as the resulting imaging signal is negative. Whilst early changes in GSH were detectable following ICI treatment, with a consequent decrease in [^18^F]FSPG, these redox changes occurred prior to immune cell influx, indicating a possible role for tumor-resident immune cells in the initial response to these immunotherapies. The cell identity and the mechanisms governing these initial redox changes still need to be elucidated. Redox homeostasis and tumor growth were restored by liproxstatin-1, indicating that ferroptotic mechanisms are at least partially responsible for the efficacy of checkpoint inhibitors. It is also known that multiple other types of cell death, such as apoptosis, necroptosis, and pyroptosis, can follow immune checkpoint blockade and that the ferroptotic response and its immunogenicity are complex and potentially context-dependent (4, 41–44). Changes in [^18^F]FSPG signal also occur during oxidative stress induced by non-ferroptotic mechanisms during chemotherapy (20) and radiotherapy (45). It is therefore important to exclude the influence of these alternative cell death pathways on the imaging signal by including ferroptosis inhibitors in the study design, whilst providing clear evidence of lipid peroxidation and the absence of markers for other forms of cell death. If properly controlled for, [^18^F]FSPG PET holds great potential as a companion diagnostic for novel ferroptosis-inducing therapies, such as bioavailable GPX4 inhibitors currently under development (46).

In conclusion, we show a direct link between [^18^F]FSPG retention, profound redox-related changes, and ferroptotic cell death. [^18^F]FSPG PET was an early non-invasive marker of immune checkpoint blockade *in vivo*. Given that [^18^F]FSPG has already been clinically translated, [^18^F]FSPG PET has great potential as a tool to aid the development of novel ferroptosis-inducing agents and in the clinic to monitor the efficacy of immunotherapies.

## Materials and methods

### Cell lines

All cell lines were grown in 1640 RPMI complete medium (10% Fetal bovine serum, ThermoFisher Scientific) supplemented with 2 mM L-glutamine (ThermoFisher Scientific) and 100 U/mL penicillin, 100 µg/mL streptomycin (Sigma Aldrich). The cells were kept at 37°C in a humidified atmosphere with 5% CO_2_. The human sarcoma cell line HT1080 and the mouse myeloma cell line B16-F0 were acquired from LGC Standards (Teddington, United Kingdom), ID8 cells were acquired from Sigma Aldrich, A549 from LGC Standards and A549 NRF2 knockout (A549 NRF2 KO) and A549 NRF2 knockout, restored (A549 NRF2 KO-R) were generously provided by Gina DeNicola. All cell lines were kept within 15-20 passages of the purchased cell stock and detached using 0.25% trypsin (Gibco). Cells were screened month for mycoplasma infection by sending of media samples to Eurofins Genomics.

### Induction and Inhibition of Ferroptosis

To induce ferroptosis in cells, the following drugs were used *in vitro*: RSL3 ((1S,3R)-Methyl 2-(2-chloroacetyl)-2,3,4,9-tetrahydro-1-[4-(methoxycarbonyl)phenyl]-1H-pyrido[3,4-b]indole-3-carboxylate) from Enzo Biochem Inc. (Farmingdale, NY, USA) (23) and iFSP1 (1-Amino-3-(4-methylphenyl)-pyrido[1,2-a]benzimidazole-2,4-dicarbonitrile) (13) from Cayman Chemicals (Ann Harbour, MI, USA). The following were used *in vivo* to introduce ferroptosis: Anti-PDL-1 antibody (Bio X Cell, Lebanon, NH, USA) and anti-CTLA4 antibody (Bio X Cell) (4). Syrian hamster IgG (Bio X Cell) and rat IgG2b, κ (Bio X Cell) were used as isogenic controls. Liproxstatin-1 (Sigma-Aldrich)(23), and ferrostatin-1 (Sigma-Aldrich) (1) were used as ferroptotic inhibitors. Other cell death inhibitors used were Z-VAD-FMK for apoptosis (Tocris Bioscience subsidiary of Bio-Techne, Minneapolis, MN, USA), bafilomycin A1 for autophagy (Tocris Bioscience), and necrostatin-2 for necroptosis (Sigma-Aldrich).

### AlamarBlue®, relative survival assay

Cells were grown to a treatment density circa 20%, in black Nunc MicroWell 96-Well plates with an optical-bottom (Thermo Fischer Scientific, Waltham, MA, USA) where incubated with medium containing RSL3 for 24 hours. Control wells contained 100 nM liproxstatin-1 in addition to the ferroptosis activators. Different concentrations of the drug were achieved by serial dilution (3.75-0.06 µM). Circa six hours before end of incubation, Resazurin, alamarBlue® (Thermo Fischer Scientific), was added, corresponding to 10% of the initial total volume, the cells were monitored as both metabolic rate and cell density influence incubation time. At the end of incubation the plates where read with a GloMax microplate reader (Promega Corp, Madison, WI, USA) using a green filter cube (Ex: 525nm, Em: 580–640 nm).

### [^18^F]FSPG synthesis

Automated radiosynthesis and quality control of [^18^F]FSPG were performed as previously reported using either the GE Fastlab^TM^ (47) or the Trasis AllInOne^TM^ (48).

### [^18^F]FSPG cell uptake studies

Cells were seeded in complete RPMI-1640 medium in 10 cm petri dishes (5 × 10^5^ for HT1080 and A549, and 3.75 × 10^5^ for ID8 and B16-F0) 24 hours before treatment with ferroptotic inducers and inhibitors. The larger format was used to ensure sufficient cells following treatment, but 6-well plates were also used, with 5 × 10^4^ cells seeded per well for ID8 and B16, and 1 × 10^5^ cells per well for A549 and HT1080, resulting in a cell density at treatment of circa 20%. Treatment was done with different concentrations (see **Supplemental Table 1**) and for 1, 3, 6 or 24 hours. Cell uptake was then conducted similar to Witney et al. (49) with the following changes: 0.185 MBq/mL [^18^F]FSPG was added, cells were collected using trypsin and pelleted by spinning them down for 4 min at 1200 *× g* and 4 °C. Cells were lysed by adding 500 µL RIPA lysis buffer (Thermo Fischer Scientific), containing 1% protease and phosphatase inhibitors (HALT, Thermo Fischer Scientific). Cell-associated radioactivity was measured in 300 µL of the cell lysate in a Wallac well counter (1282 Compugamma, LKB Wallac). Data was normalized to an [^18^F]FSPG stock solution and protein concentration using a BCA assay. For correlation of relative survival using Alamar blue and [^18^F]FSPG retention in RSL3-treated cells, AlamarBlue® was added 4 hours before sampling 100 µL for the GloMax microplate reader. This was done just before adding [^18^F]FSPG to the same wells and the plates incubated for an additional 1 h at 37 °C.

### Flow cytometric measurements of cell death and lipid peroxidation

Cells were seeded in complete RPMI-1640 medium in 10 cm petri dishes at 3.75 × 10^5^ per dish for ID8 and B16 and 5 × 10^5^ per dish for A549 and HT1080. Cells were treated with RSL3 (3.75-0.06 µM) for 1, 3, 6 or 24 hours before they were washed, trypsinised and pelleted by centrifugation (1200 *× g*, 3 min) and kept on ice throughout before analysis with a BD FACSMelody™ cell sorter (Becton, Dickinson & Company Biosciences, Franklin Lakes, NJ, USA). The cells were washed with DPBS on the plate before adding trypsin. The DPBS was saved together with the removed cell medium. The plates were incubated at 37 °C for 5 min. For cell death assessment the previously removed cell growth medium were pooled together with the detached cells to stop the trypsination, and the samples were spun down at 1200 *× g* for 4 min. For lipid peroxidation only attached cells were kept. The growth medium was the removed and the cells were washed again with ice cold Hank’s balanced salt solution (HBSS; Gibco). For cell death assessment the cells were resuspended in HBSS containing a final concentration of 5 nM SYTOX Red and incubated in the absence of light at room temperature for 15 min. For assessment of lipid peroxidation, BODIPY C11 581/591 (ThermoFisher Scientific) had been added to each dish (1.5 µM) for 20 min prior to trypsinization. Subsequently cells were spun down washed and resuspended in the same way as before at a concentration of 1 million cells per mL. The cells were then passed through a 5 µm filter before 10000-20000 cells were analyzed on the BD FACSMelody™ using the 640 nm laser and the APC filter to look at SYTOX Red and the 488 nm laser and the FITC filter to look at the oxidized product of BODIPY-c11.

### Dual assessment of cell death and [^18^F]FSPG retention in cells

HT1080 or ID8 cells seeded at ∼20% density 24 h previously were either left untreated or incubated with RSL3 (3.75 µM), with or without 100 nM liproxstatin-1, for two hours. Subsequently, 0.185 MBq [^18^F]FSPG was added to the medium of each well, and the cells were incubated for an additional 45 min at 37°C before being trypsinised, harvested by centrifugation (1200 *× g*, 3 min, 4°C), and incubated with SYTOX Red as described above. Collected cells were then sorted (∼50,000-200,000) into one negative and one SYTOX Red (640 nm laser, APC filter) positive population, using the cell sorter function on a BD FACSMelody™ (Becton, Dickinson & Company Biosciences). Unstained cells were used as reference. The two populations were then each counted in a well counter (1282 Compugamma, LKB Wallac) with data normalized to the number of cells sorted into each sample.

### Amino acid metabolomics

Extraction solvent was made the day before by mixing methanol (50%), acetonitrile (30%) and deionised water (20%) and stored overnight at -20°C. A549 and HT1080 cells, seeded as above in 10 cm dishes, were treated with 0.93 µM RSL3 alone or in combination with 100 nM liproxstatin-1 or ferrostatin-1 for 24 hours in triplicates. An untreated control was also included for calibration. To each dish, after removing and saving the media, warm PBS was used to wash the cells three times. This PBS was pooled with the removed media. Thereafter cells were trypsinised from the plate and pooled with the floating cells. The cells were then counted using a Countess™ 3 Automated Cell Counter (Thermo Fischer Scientific). After spinning down (1200 *× g*, 4 min, 4 °C) and removing the media, the cells were resuspended in ice-cold PBS. This was repeated, washing the cells three times, after which ice-cold extraction solvent was added to each cell pellet so that the final concentration was ∼4 *×* 10^6^ cells/mL. The tubes were spun at 21,000 *× g* at 1°C for 10 min. The supernatant and cell extract was retained, and the samples were stored at -80°C. 20 µL of the cell media samples were added to 480 µL of extraction buffer and mixed. Samples in extraction solvent were analyzed by LC-MS as described previously (21).

### *In vivo* animal models

Tumor growth was monitored with a digital caliper and the tumor volume was calculated using the formula for an ellipsoid. For immunotherapy studies, 1 × 10^5^ B16-F0 cells were subcutaneously injected into the right flank of female C57/BL6 mice (Charles River Laboratories, United Kingdom). Tumor burden was monitored by caliper as above and treatments were initiated when the volume reached 100 mm^3^. The mice were then treated with intraperitoneal injections of 100 µg anti-PDL-1 antibody (Bio X Cell) and 100 µg anti-CTLA4 antibody (Bio X Cell) every three days. A separate cohort of mice were additionally administered 20 mg/kg liproxstatin-1 in combination with the checkpoint inhibitors. Control mice were injected with 100 µg Syrian hamster IgG (Bio X Cell) and 100 µg Rat IgG2b, κ (Bio X Cell) isotype control antibodies according to the regimen above.

### PET/CT imaging

PET Imaging was conducted 24 hours prior to and after initiation of treatment and was repeated six to nine days later. Prior to imaging, the tail-vein was cannulated under anesthesia with 1-2.5% isoflurane in O_2_ and moved to a four-bed animal imaging hotel, which maintained the same level of anesthesia and bed temperature (37°C) throughout radiotracer administration and PET scan. Using a NanoPET/CT Plus (Mediso; Budapest, Hungary) together with a four-bed animal imaging hotel (Mediso) PET data was acquired 40 min post-injection for 20 min as previously described (18). Mice received a single bolus i.v. injection through a tail vein cannula of ∼3 MBq [^18^F]FSPG in 100 μL PBS.

### GSH

Total GSH levels were determined using cells and *ex vivo* tissues lysed in 1× passive lysis buffer. These samples were analyzed using a commercial luminescence-based kit (GSH/GSSG-Glo Assay, Promega) and a previously published method (18). In short, the assay was conducted according to the manufacturer’s instructions, with 5 µL lysate added per well of a white 96-well plate which included 5 µL of GSH standards (1-100 µmol/L). Luminescence assay readout was measured using a GloMax® plate reader (Promega). Total GSH was normalized to protein concentration, as determined by BCA assay kit (ThermoFisher Scientific).

### Western blot analysis

Western blot analysis was carried out using a previously published method (50) for antibody immunoblotting adjusted to fit the iBind Flex system (ThermoFisher Scientific) which has also been previously described (18, 20). Cells were collected as previously described for [^18^F]FSPG cell uptake studies. Both cells and frozen tissue samples were lysed using RIPA buffer containing 1× protease and phosphatase inhibitors (ThermoFisher Scientific). Rabbit monoclonal antibodies against human and murine ACSL4 (dilution, 1:200; Santa Cruz Biotechnology: sc-365230, Dallas, TX, USA), human GPX4 (dilution, 1:1000; Abcam: ab41787, Cambridge, United Kingdom), murine GPX4 (dilution, 1:1000; Abcam: ab125066), murine xCT (dilution, 1:500; Novus Biologicals: NB300-318, Bio-Techne), human xCT (dilution, 1:1000; Cell Signaling Technology: 12691, Danvers, MA, USA), human and murine NRF2 (dilution, 1:500; Cell Signaling Technology: 12721), FTH1 (dilution, 1:500; Cell Signaling Technology: 4393), NCOA4 (dilution, 1:500; Cell Signaling Technology: 66849) and FSP1/AIFM2 (dilution, 1:1000; Cell Signaling Technology: 24972) were used. Actin was used as a loading control for all experiments (anti-human and anti-mouse; dilution, 1:1000, Cell Signaling Technology; 8457), with an HRP-linked anti-rabbit IgG secondary antibody (dilution, 1:200; Cell Signaling Technology: 7074) or HRP-linked anti-mouse IgG secondary antibody (dilution, 1:200; Cell Signaling Technology: 7076). After incubation, membranes were washed five times, each wash 5 min on a rocking table, in 50 mL of tris-buffered saline with 0.1% Tween 20 (TBST). 4 mL of Amersham^™^ ECL Prime Western Blotting Detection Reagent (GE HealthCare) was added to each membrane and allowed to react for 1 min in the dark. Images of the now visualized membranes were acquired with an iBright CCD camera (Invitrogen), operated within the linear range of the camera to prevent overexposure.

### Immunohistochemical staining

B16-F0 xenografts were excised directly after sacrifice and placed in 4% Paraformaldehyde for 2-3 days. Thereafter, they were dehydrated and embedded in paraffin. Mounted successive sections of embedded tumor were then used for IHC analysis. All stainings were done using the Roche Ventana Discovery XT (Ventana Medical, Roche, Basel, Switzerland). Slides were deparaffinized in the Ventana Discovery XT platform using EZ prep solution (75°C, 8 min 950-100, Ventana Medical, Roche). Tris-EDTA buffer (95°C, 32 min, 950-300, Ventana Medical, Roche) was used for antigen retrieval and blocking of endogenous peroxidases was done on the Ventana Discovery XT using Inhibitor CM (37°C, 4 min, Ventana Medical, Roche). Slides were thereafter incubated with anti-mouse CD8 antibody (1 hour, 1:100; ab217344, Abcam), anti-mouse F4/80 (4 hours, 1:100 dilution; 70076, Cell Signaling), anti-4-HNE (1 hour, 1:100 dilution; ab48506, Abcam), anti-Ki67 (4 hours, 1:100 dilution; 12202, Cell Signaling), anti-mouse CD4 (1 hour, 1:100 dilution; ab183685, Abcam), or anti-cleaved caspase three antibodies (32 min, 1:100 dilution; 9661, Cell Signaling). For all but 4-HNE, a 1:200 dilution of secondary antibody (goat anti-rabbit; ab207995-500 µg, Abcam) was used (incubation: Caspase-3, 44 min; CD4 and CD8, 32 min; F4:80, 1 hour). For 4-HNE, a 1 hour incubation with a 1:200 dilution of secondary antibody (rabbit anti-mouse; ab98668, Abcam) was performed. After washing on the Ventana Discovery XT platform, sections were incubated with DAB substrate solution (760-124, Ventana Medical, Roche) and counterstained using Roche counterstain kits (Haematoxylin 760-2021/Bluing Reagent 760-2037). The slides were then washed in warm tap water and dehydrated in graded ethanol and xylene. Using permanent mounting media (Pertex, 00811, Histolab), the slides were attached to a coverslip, and images were later acquired using a NanoZoomer (Hamamatsu).

### *Ex vivo* analysis of lipid peroxidation

Lipid peroxidation in frozen tissue samples was assessed using an MDA kit (ab118970, Abcam) according to the manufacturer’s instructions. In short, MD lysis buffer and butylated hydroxytoluene stock solution was brought to room temperature mixed and added to Lysing Matrix tubes with a total of 300 µL lysis solution per sample. Thawed tissue, kept on ice, was washed with ice cold PBS and then lysed at 4°C, as described above, thereafter centrifuged at 13,000 × *g* for 10 min and 4°C to remove debris and collect the supernatant. A Pierce BCA assay was performed on all supernatants, as described above to determine protein concentration. MDA standards were made each time by diluting the 4.17 M stock solution appropriately and according to instructions with ddH_2_O. Developer solution was also made new each time with a new vial of thiobarbituric acid reconstituted in 7.5 mL glacial acetic acid. The slurry was then moved to a new tube and 25 mL of ddH_2_O was added (the solution was sonicated if needed). To each sample, 0.6 mL of developer solution was added and the samples were then incubated at 95°C for 60 min. From each sample, 0.2 mL was moved to a 96-well plate to determine the optical absorbance at 532 nm using a Varioskan plate reader (ThermoFisher Scientific) and MDA quantified using the standard curve.

### Data analysis

All flow cytometry data were analyzed using FlowJo v10.6.2 (Becton, Dickinson & Company). All final data were expressed as the mean ± one standard deviation (SD) except for fluorescence, which was expressed as the median fluorescence intensity ± one standard deviation (SD). GraphPad Prism (version 8.4.2, GraphPad Software, LCC) was used for statistical analysis, linear regression and non-linear curve fitting, and IC_50_ calculations. For statistical significance across multiple samples, 1-way analysis of variance (ANOVA) followed by T-tests multiple comparison correction with adjusted P-values (Tukey’s or Dunnett’s method) were performed. *In vitro* data were acquired on separate days in at least three biological repeats. The reconstructed PET data was analyzed using 3D regions of interest (ROI) comprising of manually and sequentially drawing 2D regions on the CT images of the xenografts using VivoQuant software (v.2.5, Invicro Ltd.). The quantified radioactivity present in each ROI was expressed as a percentage of the injected dose per gram of tissue volume (% ID/g). Immunohistochemical staining was analyzed using QuPath (v0.50) (51). For stained tissue sections from B16-F0 xenografts treated with immune checkpoint inhibitors or isotype control antibodies, one to three areas of each tissue section were randomly selected. Areas rich in melanin were excluded to facilitate the analysis. ROIs were then created for tumor-containing regions. Thereafter, the total number of cells and the number of positively stained cells were detected using the positive cell detection tool in QuPath, and the fraction of positive cells was calculated.

## Acknowledgements

We would like to thank Prof. Gina de Nicola for providing the A549 NRF2 knockout cell line. At King’s College London, we would like to extend our gratitude to Jana Kim and Kavitha Sunassee for their help with preclinical imaging and in vivo protocols and Home Office Licence, as well as the technical team at BMEIS: Matthew Hutchings, Lisa Sanderson, Shivam Bhardwaj, Louis Kitchenham, and David Thakor for their work. We also thank IQPath at University College of London for sectioning and immunohistochemically staining embedded xenografts. We would further like to thank Mrs. Berta Kamprad’s Cancer Foundation for supporting Oskar Vilhelmsson Timmermand during the drafting of the manuscript.

## Author Contribution Statement

THW conceived the study and acquired funding. THW and OVT designed the study and wrote the manuscript. OVT, ARB, MEG, SNS, HEG, RSE, MT, AHU, WET, JB and RF performed the experiments and analyzed the data. OM provided metabolomic data interpretation. NP, MMM and JvdA contributed to the discussion and data interpretation. All authors read and approved the final manuscript.

## Ethics Statement

All animal experiments were performed in accordance with the United Kingdom Home Office Animal (Scientific Procedures) Act 1986 under project license PP9982297 and received local Animal Welfare and Ethical Review Body approval.

## Data Availability Statement

The original contributions presented in the study are included in the main text or the supplementary materials. Other data that support the findings are available from the corresponding author upon reasonable request.

## References

1. Dixon SJ, Lemberg KM, Lamprecht MR, Skouta R, Zaitsev EM, Gleason CE, et al. Ferroptosis: an iron-dependent form of nonapoptotic cell death. Cell. 2012;149(5):1060–72.

2. Stockwell BR, Friedmann Angeli JP, Bayir H, Bush AI, Conrad M, Dixon SJ, et al. Ferroptosis: A Regulated Cell Death Nexus Linking Metabolism, Redox Biology, and Disease. Cell. 2017;171(2):273–85.

3. Conrad M, Kagan VE, Bayir H, Pagnussat GC, Head B, Traber MG, et al. Regulation of lipid peroxidation and ferroptosis in diverse species. Genes Dev. 2018;32(9-10):602–19.

4. Wang W, Green M, Choi JE, Gijon M, Kennedy PD, Johnson JK, et al. CD8(+) T cells regulate tumour ferroptosis during cancer immunotherapy. Nature. 2019;569(7755):270–4.

5. Lei G, Zhang Y, Koppula P, Liu X, Zhang J, Lin SH, et al. The role of ferroptosis in ionizing radiation-induced cell death and tumor suppression. Cell Res. 2020;30(2):146–62.

6. Lang X, Green MD, Wang W, Yu J, Choi JE, Jiang L, et al. Radiotherapy and Immunotherapy Promote Tumoral Lipid Oxidation and Ferroptosis via Synergistic Repression of SLC7A11. Cancer Discov. 2019;9(12):1673–85.

7. Li Y, Feng D, Wang Z, Zhao Y, Sun R, Tian D, et al. Ischemia-induced ACSL4 activation contributes to ferroptosis-mediated tissue injury in intestinal ischemia/reperfusion. Cell Death Differ. 2019;26(11):2284–99.

8. Chen Y, Fang ZM, Yi X, Wei X, Jiang DS. The interaction between ferroptosis and inflammatory signaling pathways. Cell Death Dis. 2023;14(3):205.

9. Mishima E, Nakamura T, Doll S, Proneth B, Fedorova M, Pratt DA, et al. Recommendations for robust and reproducible research on ferroptosis. Nat Rev Mol Cell Biol. 2025;26(8):615–30.

10. Doll S, Proneth B, Tyurina YY, Panzilius E, Kobayashi S, Ingold I, et al. ACSL4 dictates ferroptosis sensitivity by shaping cellular lipid composition. Nat Chem Biol. 2017;13(1):91–8.

11. Kagan VE, Mao G, Qu F, Angeli JP, Doll S, Croix CS, et al. Oxidized arachidonic and adrenic PEs navigate cells to ferroptosis. Nat Chem Biol. 2017;13(1):81–90.

12. Murphy TH, Miyamoto M, Sastre A, Schnaar RL, Coyle JT. Glutamate toxicity in a neuronal cell line involves inhibition of cystine transport leading to oxidative stress. Neuron. 1989;2(6):1547–58.

13. Doll S, Freitas FP, Shah R, Aldrovandi M, da Silva MC, Ingold I, et al. FSP1 is a glutathione-independent ferroptosis suppressor. Nature. 2019;575(7784):693–8.

14. Yang WS, SriRamaratnam R, Welsch ME, Shimada K, Skouta R, Viswanathan VS, et al. Regulation of ferroptotic cancer cell death by GPX4. Cell. 2014;156(1-2):317–31.

15. Ursini F, Maiorino M, Valente M, Ferri L, Gregolin C. Purification from pig liver of a protein which protects liposomes and biomembranes from peroxidative degradation and exhibits glutathione peroxidase activity on phosphatidylcholine hydroperoxides. Biochim Biophys Acta. 1982;710(2):197–211.

16. Atkuri KR, Mantovani JJ, Herzenberg LA, Herzenberg LA. N-Acetylcysteine--a safe antidote for cysteine/glutathione deficiency. Curr Opin Pharmacol. 2007;7(4):355–9.

17. Stockwell BR, Jiang X. The Chemistry and Biology of Ferroptosis. Cell Chem Biol. 2020;27(4):365–75.

18. Greenwood HE, Barber AR, Edwards RS, Tyrrell WE, George ME, Dos Santos SN, et al. Imaging NRF2 activation in non-small cell lung cancer with positron emission tomography. Nat Commun. 2024;15(1):10484.

19. Anandhan A, Dodson M, Schmidlin CJ, Liu P, Zhang DD. Breakdown of an Ironclad Defense System: The Critical Role of NRF2 in Mediating Ferroptosis. Cell Chem Biol. 2020;27(4):436–47.

20. McCormick PN, Greenwood HE, Glaser M, Maddocks ODK, Gendron T, Sander K, et al. Assessment of Tumor Redox Status through (S)-4-(3-[(18)F]fluoropropyl)-L-Glutamic Acid PET Imaging of System x(c) (-) Activity. Cancer Res. 2019;79(4):853–63.

21. Greenwood HE, McCormick PN, Gendron T, Glaser M, Pereira R, Maddocks ODK, et al. Measurement of Tumor Antioxidant Capacity and Prediction of Chemotherapy Resistance in Preclinical Models of Ovarian Cancer by Positron Emission Tomography. Clin Cancer Res. 2019;25(8):2471–82.

22. Smith LM, Greenwood HE, Tyrrell WE, Edwards RS, de Santis V, Baark F, et al. The chicken chorioallantoic membrane as a low-cost, high-throughput model for cancer imaging. Npj Imaging. 2023;1(1):1.

23. Yang WS, Stockwell BR. Synthetic lethal screening identifies compounds activating iron-dependent, nonapoptotic cell death in oncogenic-RAS-harboring cancer cells. Chem Biol. 2008;15(3):234–45.

24. Parker JL, Deme JC, Kolokouris D, Kuteyi G, Biggin PC, Lea SM, et al. Molecular basis for redox control by the human cystine/glutamate antiporter system xc(). Nat Commun. 2021;12(1):7147.

25. Zhang DD. Thirty years of NRF2: advances and therapeutic challenges. Nat Rev Drug Discov. 2025.

26. Torrente L, DeNicola GM. Targeting NRF2 and Its Downstream Processes: Opportunities and Challenges. Annu Rev Pharmacol Toxicol. 2022;62:279–300.

27. Dodson M, Castro-Portuguez R, Zhang DD. NRF2 plays a critical role in mitigating lipid peroxidation and ferroptosis. Redox Biol. 2019;23:101107.

28. Kang YP, Torrente L, Falzone A, Elkins CM, Liu M, Asara JM, et al. Cysteine dioxygenase 1 is a metabolic liability for non-small cell lung cancer. Elife. 2019;8.

29. Dai E, Zhang W, Cong D, Kang R, Wang J, Tang D. AIFM2 blocks ferroptosis independent of ubiquinol metabolism. Biochem Biophys Res Commun. 2020;523(4):966–71.

30. Koppula P, Lei G, Zhang Y, Yan Y, Mao C, Kondiparthi L, et al. A targetable CoQ-FSP1 axis drives ferroptosis- and radiation-resistance in KEAP1 inactive lung cancers. Nat Commun. 2022;13(1):2206.

31. Mishima E, Nakamura T, Doll S, Proneth B, Fedorova M, Pratt DA, et al. Recommendations for robust and reproducible research on ferroptosis. Nat Rev Mol Cell Biol. 2025.

32. Sonay AY, Apfelbaum E, Larney BEM, Grimm J. Phosphatidylserine exposure and plasma membrane perforation as ferroptotic signatures for in vivo imaging. Npj Imaging. 2025;3(1):48.

33. Zeng F, Nijiati S, Liu Y, Yang Q, Liu X, Zhang Q, et al. Ferroptosis MRI for early detection of anticancer drug-induced acute cardiac/kidney injuries. Sci Adv. 2023;9(10):eadd8539.

34. Colovic M, Yang H, Southcott L, Merkens H, Colpo N, Benard F, et al. Comparative Evaluation of [(18)F]5-Fluoroaminosuberic Acid and (4S)-4-3-[(18)F]fluoropropyl)-l-Glutamate as System xC--Targeting Radiopharmaceuticals. J Nucl Med. 2023;64(8):1314–21.

35. Greenwood HE, Edwards R, Koglin N, Berndt M, Baark F, Kim J, et al. Radiotracer stereochemistry affects substrate affinity and kinetics for improved imaging of system x(C)(-) in tumors. Theranostics. 2022;12(4):1921–36.

36. Moses A, Malek R, Kendirli MT, Cheung P, Landry M, Herrera-Barrera M, et al. Monitoring of cancer ferroptosis with [(18)F]hGTS13, a system xc- specific radiotracer. Theranostics. 2025;15(3):836–49.

37. Caneque T, Baron L, Muller S, Carmona A, Colombeau L, Versini A, et al. Activation of lysosomal iron triggers ferroptosis in cancer. Nature. 2025.

38. Liao P, Wang W, Wang W, Kryczek I, Li X, Bian Y, et al. CD8(+) T cells and fatty acids orchestrate tumor ferroptosis and immunity via ACSL4. Cancer Cell. 2022;40(4):365–78 e6.

39. Bensch F, van der Veen EL, Lub-de Hooge MN, Jorritsma-Smit A, Boellaard R, Kok IC, et al. (89)Zr-atezolizumab imaging as a non-invasive approach to assess clinical response to PD-L1 blockade in cancer. Nat Med. 2018;24(12):1852–8.

40. Farwell MD, Gamache RF, Babazada H, Hellmann MD, Harding JJ, Korn R, et al. CD8-Targeted PET Imaging of Tumor-Infiltrating T Cells in Patients with Cancer: A Phase I First-in-Humans Study of (89)Zr-Df-IAB22M2C, a Radiolabeled Anti-CD8 Minibody. J Nucl Med. 2022;63(5):720–6.

41. Fan F, Liu P, Bao R, Chen J, Zhou M, Mo Z, et al. A Dual PI3K/HDAC Inhibitor Induces Immunogenic Ferroptosis to Potentiate Cancer Immune Checkpoint Therapy. Cancer Res. 2021;81(24):6233–45.

42. Yang F, Xiao Y, Ding JH, Jin X, Ma D, Li DQ, et al. Ferroptosis heterogeneity in triple-negative breast cancer reveals an innovative immunotherapy combination strategy. Cell Metab. 2023;35(1):84–100 e8.

43. Drijvers JM, Gillis JE, Muijlwijk T, Nguyen TH, Gaudiano EF, Harris IS, et al. Pharmacologic Screening Identifies Metabolic Vulnerabilities of CD8(+) T Cells. Cancer Immunol Res. 2021;9(2):184–99.

44. Liu T, Zhu C, Chen X, Guan G, Zou C, Shen S, et al. Ferroptosis, as the most enriched programmed cell death process in glioma, induces immunosuppression and immunotherapy resistance. Neuro Oncol. 2022;24(7):1113–25.

45. Sambasivan K, Tyrrell WE, Farooq R, Mynerich J, Edwards RS, Tanc M, et al. [(18)F]FSPG-PET provides an early marker of radiotherapy response in head and neck squamous cell cancer. Npj Imaging. 2024;2(1):28.

46. Rodencal J, Kim N, He A, Li VL, Lange M, He J, et al. Sensitization of cancer cells to ferroptosis coincident with cell cycle arrest. Cell Chem Biol. 2024;31(2):234–48 e13.

47. Edwards R, Greenwood HE, McRobbie G, Khan I, Witney TH. Robust and Facile Automated Radiosynthesis of [(18)F]FSPG on the GE FASTlab. Mol Imaging Biol. 2021;23(6):854–64.

48. Farooq R, Gendron T, Edwards RS, Witney TH. Compact and cGMP-compliant automated synthesis of [(18)F]FSPG on the Trasis AllinOne. EJNMMI Radiopharm Chem. 2025;10(1):2.

49. Witney TH, Fortt R, Aboagye EO. Preclinical assessment of carboplatin treatment efficacy in lung cancer by 18F-ICMT-11-positron emission tomography. PLoS One. 2014;9(3):e91694.

50. Witney TH, Kettunen MI, Day SE, Hu DE, Neves AA, Gallagher FA, et al. A comparison between radiolabeled fluorodeoxyglucose uptake and hyperpolarized (13)C-labeled pyruvate utilization as methods for detecting tumor response to treatment. Neoplasia. 2009;11(6):574–82, 1 p following 82.

51. Bankhead P, Loughrey MB, Fernandez JA, Dombrowski Y, McArt DG, Dunne PD, et al. QuPath: Open source software for digital pathology image analysis. Sci Rep. 2017;7(1):16878.

