## Supplementary data for "System x_c_^−^ imaging maps ferroptosis-linked redox remodeling in cancer"

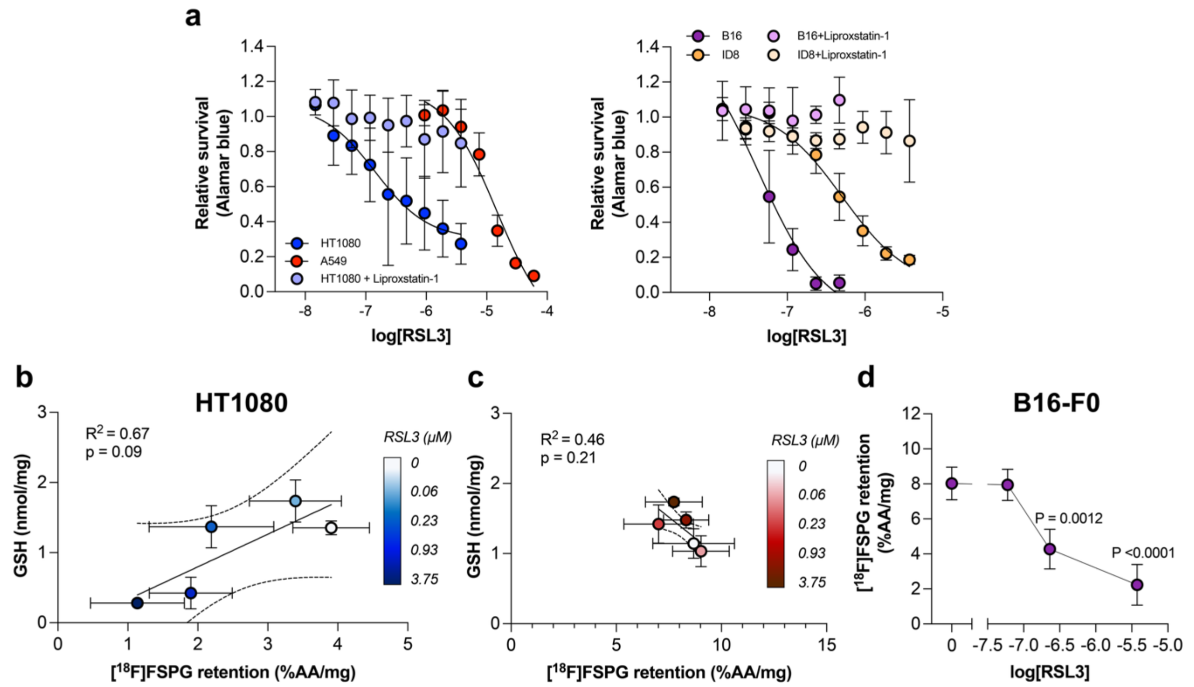


***Supplementary Figure 1. RSL3 treatment induces cell death, depletes GSH, and reduces [$^{18}$F]FSPG retention in sensitive but not resistant cells. a.*** *The relative survival of HT1080, A549, B16 and ID8 following treatment with RSL3 at increasing concentrations (0-60 µM) for 24 h. Cell survival was evaluated with AlamarBlue® following RSL3 treatment, with and without 100 nM of liproxstatin-1.* ***b.*** *Correlation plot of GSH versus [$^{18}$F]FSPG retention in HT1080 cells following RSL3 treatment. Dotted lines represent the 95% confidence interval.* ***c.*** *Correlation plot of GSH versus [$^{18}$F]FSPG retention in A549 cells. Dotted lines represent the 95% confidence interval.* ***d.*** *Changes in [ $^{18}$F]FSPG retention following increasing concentrations of RSL3 in the B16-F0 cells.*

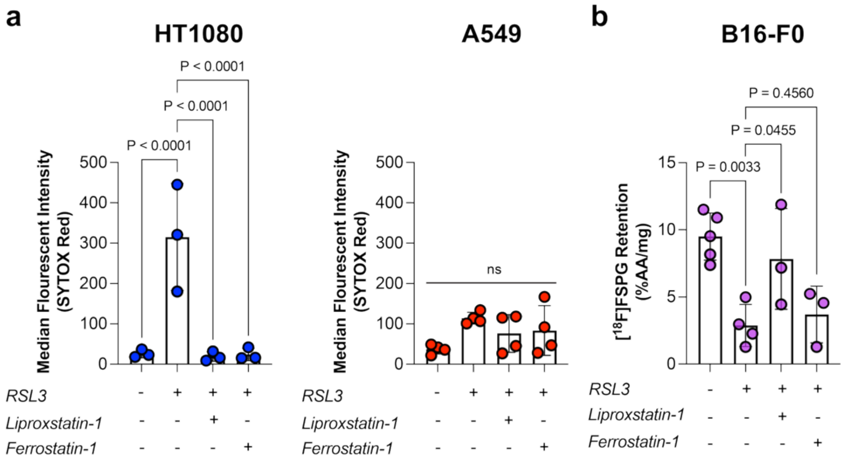


***Supplementary Fig. 2. Ferroptosis inhibitors rescue cell death and [$^{18}$F]FSPG retention in sensitive cells following RSL3 treatment. a.*** *Necrotic phenotype, characterized by Sytox Red staining in HT1080 and A549 cells treated with RSL3 only and in combination with ferroptosis inhibitors.* ***b.*** *[$^{18}$F]FSPG retention in B16-F0 cells treated with RSL3 only or in combination with ferroptosis inhibitors. ns, not significant.*

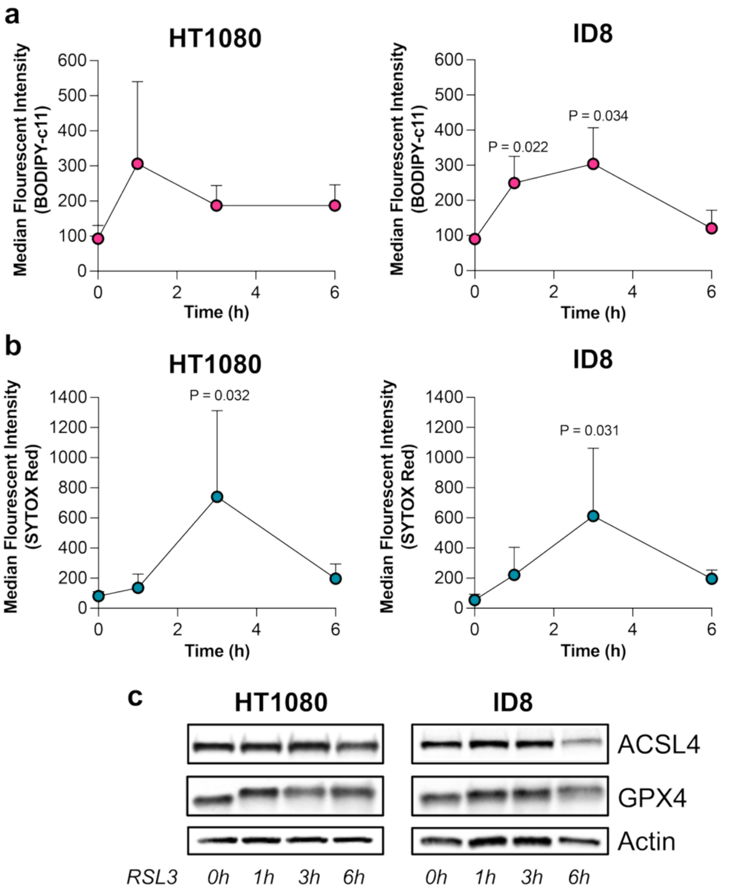


***Supplementary Figure 3. Temporal changes following RSL3 treatment in HT1080 and ID8 cells.*** ***a.*** *Lipid peroxidation in HT1080 and ID8 cells 0-6 h after treatment with 3.75 µM.* ***b.*** *RSL3-induced necrosis in HT1080 and ID8 cells 0-6 h after treatment, measured with SytoxRed.* ***c****. Protein expression of ACSL4 and GPX4 in HT1080 and ID8 cells treated between 0-6 h with 3.75 µM RSL3. Actin was used as a loading control.*

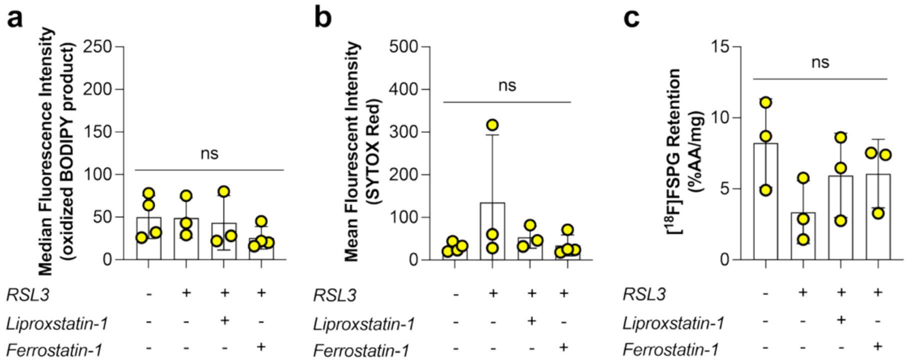


***Supplementary Figure 4. Ectopic NRF2 expression in NRF2 KO A549 cells restores ferroptosis resistance. a-c.*** *Lipid peroxidation (**a**), necrosis (**b**) and [$^{18}$F]FSPG retention (**c**) in A549 NRF2 knockout NRF2-restored cells treated with RSL3 (0.93 µM) with/without co-treatment with liproxstatin-1 (100 nM) or ferrostatin-1 (12 µM) for 24 h and compared to untreated cells (DMSO).*

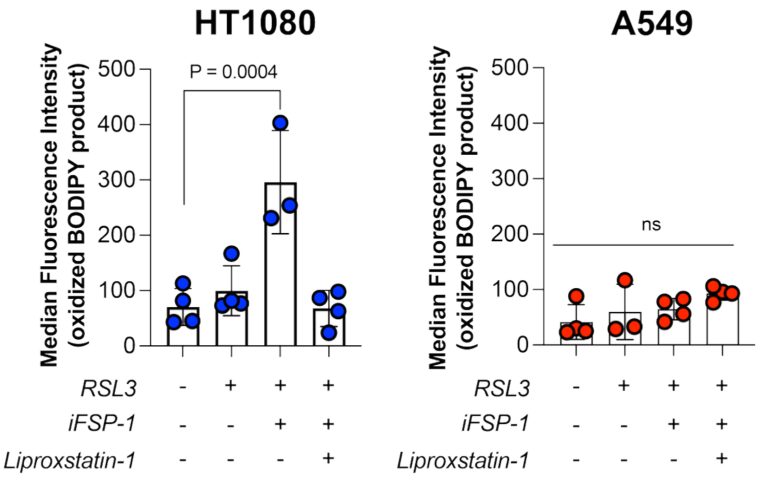


***Supplementary Figure 5. Lipid peroxidation in HT1080 and A549 cells following treatment with a sub-effective dose of RSL3 in the presence and absence of iFSP-1.***

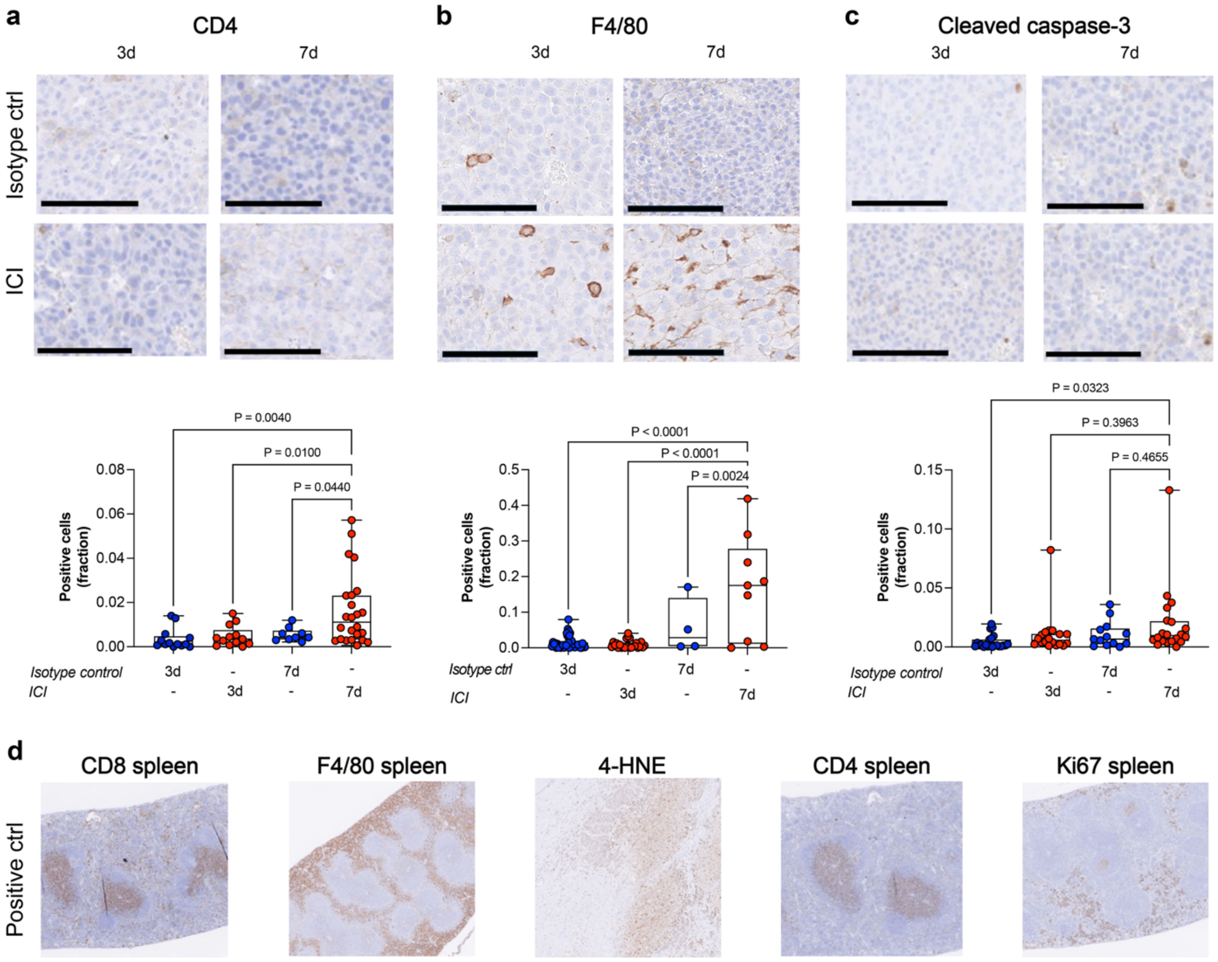


***Supplementary Figure 6. CD4, F4/80 and cleaved caspase-3 staining following immune checkpoint blockade. a-c.*** *Representative images of immunohistochemical staining and semi-quantitative analysis of the fraction of stained cells following immune checkpoint blockade for CD4 (**a**), F4/80 (**b**) and cleaved caspase-3 (**c**). Scale bar, 100 µm.* ***d.*** *Positive control staining in spleen for CD8, F4/80, CD4 and Ki67, and 4-HNE in oocytes confirming that the immunohistochemical procedure is working as intended.*

**Supplementary Table 1. Concentrations used for RSL3 treatment**

| Cell line | [RSL3] (μM) | | | |
|---|---|---|---|---|
| | 24-hour incubation | With cell death inhibitors | 1-, 3- and 6-hour time course | With iFSP-1 |
| HT1080 | 3.75, 0.93, 0.23, 0.06 | 0.93 | 3.75 | 0.06 |
| A549 | 3.75, 0.93, 0.23, 0.06 | 0.93 | - | 0.93 |
| B16-F0 | 3.75, 0.23, 0.06 | 0.23 | - | - |
| ID8 | - | - | 3.75 | - |
| A549 NRF2 KO | - | 0.93 | - | - |
| A549 NRF2 KO-R | - | 0.93 | - | - |
